# A deep learning approach for staging embryonic tissue isolates with small data

**DOI:** 10.1101/2020.07.15.204735

**Authors:** Adam Pond, Seongwon Hwang, Berta Verd, Benjamin Steventon

**Affiliations:** Department of Genetics, University of Cambridge, Cambridge CB2 3EH, UK; Department of Zoology, University of Oxford, Oxford OX1 3SZ, UK

## Abstract

Machine learning approaches are becoming increasingly widespread and are now present in most areas of research. Their recent surge can be explained in part due to our ability to generate and store enormous amounts of data with which to train these models. The requirement for large training sets is also responsible for limiting further potential applications of machine learning, particularly in fields where data tend to be scarce such as developmental biology. However, recent research seems to indicate that machine learning and Big Data can sometimes be decoupled to train models with modest amounts of data. In this work we set out to train a CNN-based classifier to stage zebrafish tail buds at four different stages of development using small information-rich data sets. Our results show that two and three dimensional convolutional neural networks can be trained to stage developing zebrafish tail buds based on both morphological and gene expression confocal microscopy images, achieving in each case up to 100% test accuracy scores. Importantly, we show that high accuracy can be achieved with data set sizes of under 100 images, much smaller than the typical training set size for a convolutional neural net. Furthermore, our classifier shows that it is possible to stage isolated embryonic structures without the need to refer to classic developmental landmarks in the whole embryo, which will be particularly useful to stage 3D culture in vitro systems such as organoids. We hope that this work will provide a proof of principle that will help dispel the myth that large data set sizes are always required to train CNNs, and encourage researchers in fields where data are scarce to also apply ML approaches.

**Author summary:** The application of machine learning approaches currently hinges on the availability of large data sets to train the models with. However, recent research has shown that large data sets might not always be required. In this work we set out to see whether we could use small confocal microscopy image data sets to train a convolutional neural network (CNN) to stage zebrafish tail buds at four different stages in their development. We found that high test accuracies can be achieved with data set sizes of under 100 images, much smaller than the typical training set size for a CNN. This work also shows that we can robustly stage the embryonic development of isolated structures, without the need to refer back to landmarks in the tail bud. This constitutes an important methodological advance for staging organoids and other 3D culture in vitro systems. This work proves that prohibitively large data sets are not always required to train CNNs, and we hope will encourage others to apply the power of machine learning to their areas of study even if data are scarce.

## Introduction

Machine learning (ML) approaches are not new, with early works dating as far back as the 1950s [1]. However, in the last two decades, the field has experienced an astonishing surge in productivity and progress. This soar can be explained, at least in part, by our new-found ability to generate and store ever larger amounts of data (Big Data) with which to train ML models coupled with unprecedented computational speed and power. It is becoming difficult to find a field that hasn’t yet adopted ML approaches in one way or another, with applications ranging all the way from the humanities (see [2–4] for some examples) to paleontology [5] or health care [6]. The biological sciences are indeed no exception [7–10], having entered into the era of Big Data thanks to technological advances such as next generation sequencing and multi-omics approaches [11]. Machine learning, and deep learning in particular, have been used to predict sequence specifities of both DNA- and RNA-binding proteins, as well as enhancers and other types of regulatory regions [12, 13]. A few other notable examples include the wide use of ML approaches to address questions in population and evolutionary genetics [14], predicting transcript abundance from DNA sequence [15], predicting methylation states in single cells [16], imputation of missing SNPs [17] as well as variant calling [18].

Deep learning, and in particular convolutional neural networks (CNNs), are being increasingly applied to problems involving various types of microscopy images in both biology and medicine [19–22]. CNNs are a family of algorithms that have gained popularity in recent years thanks to their remarkable levels of accuracy when applied to computer vision problems such as image classification, analysis and object detection [23]. The design of CNNs has been inspired by animal nervous systems: they are made up of high numbers of interconnected computational nodes (known as neurons) across various different layers, and together they manage to optimise the network parameters by learning collectively to improve the network’s performance during training [24]. CNNs differ from other machine learning algorithms also used in computer vision such as support vector machines and random forests in that they don’t rely on the manual annotation of identifiable features in the images used. Instead, CNNs take image pixels as inputs directly, without the need for any manual annotation of the images themselves. This aspect is particularly attractive for the analysis of image-based biological data as it allows for conclusions to be drawn from the image data itself, without requiring any initial image analysis, eliminating the any biasing during feature detection. CNNs have been optimised to make the most of recent advances in GPU technologies resulting in impressively high accuracies and classification speeds when compared to similar methods [25–28]. In addition, formulating and training CNNs has become more accessible in recent years thanks to the development of various user-friendly softwares [29–31].

Deep learning algorithms have been proving their prowess at classifying natural images ever since the convolutional neural network AlexNet won the ImageNet Large Scale Visual Recognition in 2012 [32]. Since then, this methodology has been successfully applied to object detection and classification in biological microscopy image data revolutionising the field. In one of the first examples, a CNN model was trained to identify and stage cells in brightfield microscopy images of malaria-infected blood [33]. Subsequently, CNNs have been used to obtain morphological profiles of cultured cells from fluorescence microscopy images [34, 35], identify changes in cell state [36–38], determine protein localisation within the cell by classifying spatial patterns in fluorescence images [39–42] and detect bacteria in a zebrafishes digestive tract [25], to name just a few examples.

While the availability of Big Data has allowed for ML approaches to be readily applied to many areas in biology, the requirement for large training sets is also responsible for limiting further potential applications. This challenge is not unique to biology. Personalised medicine, where the aim is to use data from single patients to train models that will inform their diagnosis and treatment, is another promising field which suffers from data scarcity. To overcome such pitfalls, an active area of research has emerged to find ways to train accurate ML models with smaller data set sizes to hopefully uncouple ML from Big Data [43–48].

Data in developmental biology is often scarce: embryos are rarely available in high numbers, and even when they are, staining and imaging is time consuming, limiting the amount of data that can be collected. However scarce, such data sets are also increasingly information-rich, often consisting of stacks of high-resolution z-slices that together make up a complete 3D image of the embryo or region under study. Moreover, advances in the multi-plexed detection of mRNA distribution means that such 3D structural information can be coupled with information about the distribution and quantification of multiple mRNA species from the tissue to sub-cellular level [49–52]. With ML approaches it is often unclear at the onset, exactly how much data is going to be required to achieve a certain test accuracy. Inspired by previous work showing that CNN-based classifiers could be trained with information-rich small data sets [43–48], we set out to see if we would be able to train a CNN to classify zebrafish tail buds of different stages with the limited data that was available to us in the lab.

Zebrafish (Danio rerio) is a very popular model organism in developmental biology. Its embryos go from fertilization to free swimming larvae in three days [53]. In the first day of their development, zebrafish embryos enter the segmentation period [53]. During this time, the embryo elongates, somites (precursors of the dermis, skeletal muscle, cartilage, tendons, and vertebrae) appear sequentially in an anterior to posterior manner and the tail bud becomes more prominent (Fig 1A). Zebrafish embryos are routinely staged based on their overall shape and the total number of somites that they have formed. However, during these stages the overall shape of the tail bud also changes, although more subtly, to become shorter, thinner and overall straighter (Fig 1B).

**Fig 1.**
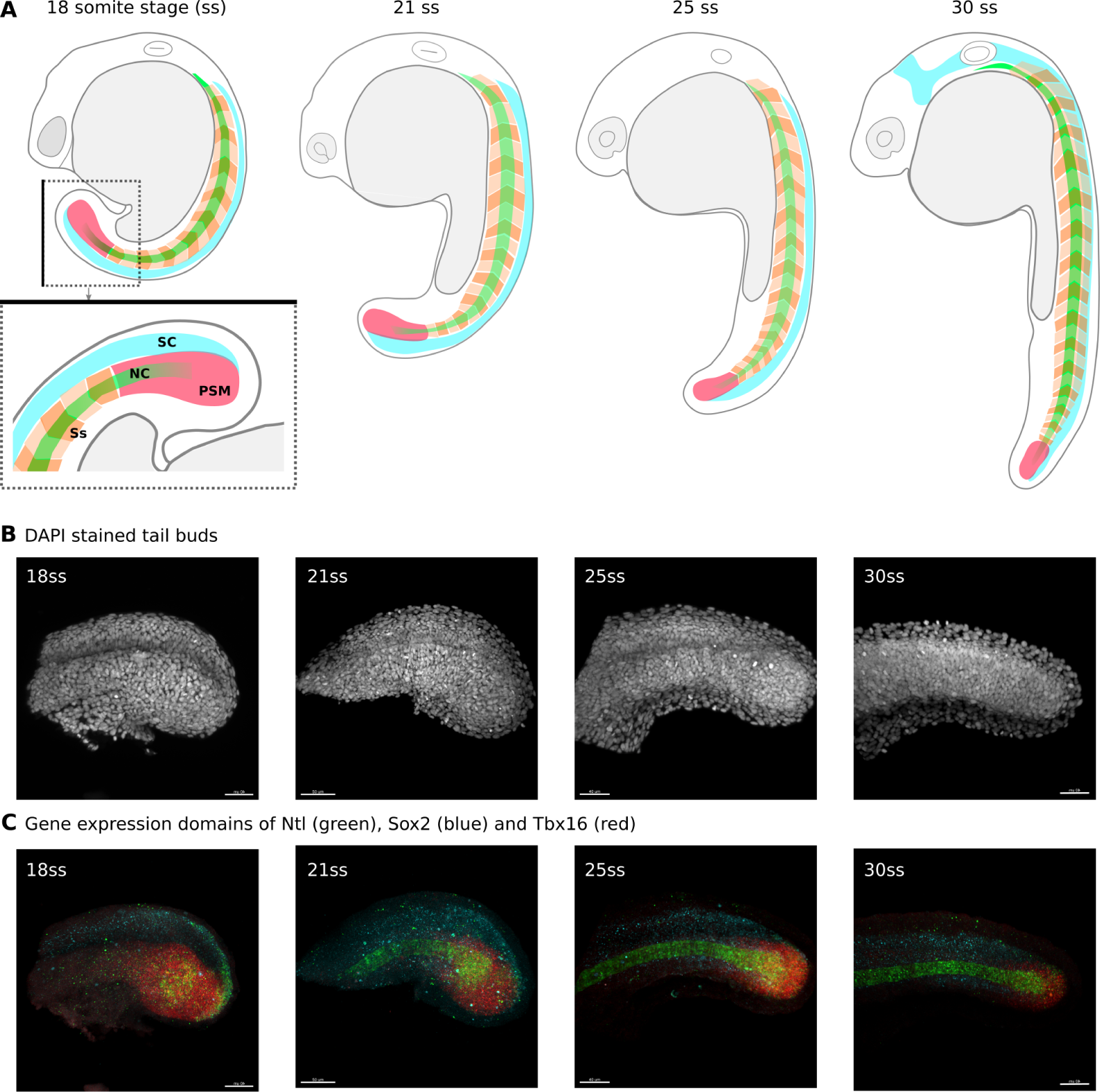
Zebrafish development and the tail bud classification problem. **A.** Schematic drawings of zebrafish embryos at 18, 21, 25 and 30 somite stages. The tail bud region of the 18 somite stage embryo schematic is shown inside the dotted square and below, rotated 90° to the right and zoomed in. Black line on the boxes to help visualise alignment. Posterior to the right, anterior to the left, dorsal up and ventral down. The spinal cord (SC) is shown in cyan, alternate somites (Ss) are shown in different shades of orange, the notochord (NC) is shown in green and the pre-somitic mesoderm (PSM) is shown in pink. **B.** Maximum projection images of the tail buds of embryos at the 18, 21, 25 and 30 somite stages respectively, stained with Dapi and imaged on a confocal microscope. **C.** Maximum projection images of the same tail buds as in B, stained for the mRNA of tbxta (green in the posterios PSM and notochord), tbx16 (red in the PSM) and sox2 (blue in the spinal cord) using HCR V.3 and imaged on a confocal microscope.

In this work we set out to test whether it would be possible to use a small albeit information-rich data set of confocal images to train a CNN to accurately classify images of zebrafish tail buds at four different stages during the segmentation period. In addition to the challenge regarding the small size of our training set, we also wanted to see whether a CNN would be able to learn to classify based on subtle changes in the shape of an isolated embryonic structure, in this case the tail bud. Such a classifier would solve the problem of asynchronous development within clutches and save man-hours by automating the staging step in laboratories across the world.

In this paper we show that, contrary to popular belief, small information-rich data sets can also be used to train CNN-based classifiers to a high accuracy. We have focused on building and training CNNs to correctly stage two and three dimensional morphological and gene expression image data of zebrafish tail buds at four different stages during the segmentation period of development. We found that CNN-based classifiers can yield test accuracies of 100% when trained with less than 100 images. Furthermore, our results show that this is the case both when morphological and gene expression image data were used as the training set. Surprisingly, higher dimensional data (3D versus 2D) isn’t always associated with a higher accuracy score. We hope that our work will provide a precedent and encourage others in the life sciences to apply ML approaches even when their data are relatively scarce.

## Results and Discussion

### 0.1 Data

In this work, we set out to train convolutional neural networks to classify 2D and 3D confocal images of dissected tailbuds taken from zebrafish embryos at four close but different stages in development: 16-18 somite stage, 20-22 somite stage, 24-26 somite stage and 28-30 somite stage (Fig 1A). The chosen classes cover approximately 1.5hrs of embryonic development each, and are fine enough for our general research purposes, which aim to understanding fate specification in the tail bud during the segmentation period.

Whole embryos were stained for three gene products, Tbxta, Tbx16 and Sox2, using HCR V.3 [54]. Tbxta is expressed in the notochord and mesoderm progenitor zone (Fig 2A and D), Tbx16 is also expressed in the mesoderm progenitor zone and is present in the posterior pre-somitic mesoderm, but not in the notochord (Fig 2A and C) [55, 56]. Sox 2 is a neural marker which is expressed in the neural progenitor zone (Fig 1C and Fig 2A and B) [57]. All embryos were also stained with DAPI, a nuclear marker (Fig 1B) which allows us to visualise the morphology of the entire tail. The staining protocol takes three days and has been optimised to allow us to stain larger numbers of embryos at a time (see Materials and Methods).

**Fig 2.**
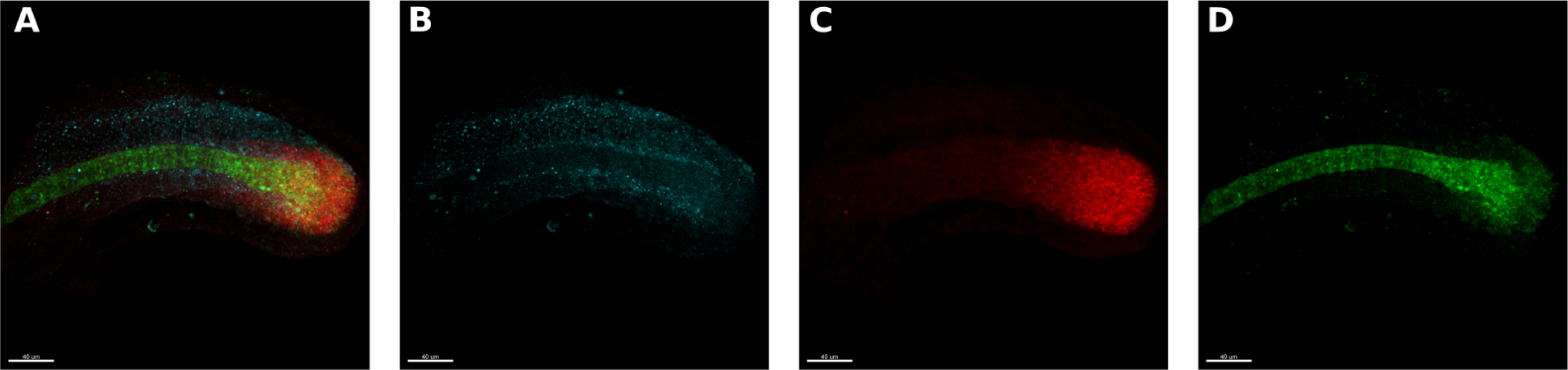
Gene expression data. **A.** Same maximum projection image as in Fig 1C (third from the left) of a 25 somite stage embryo’s tail bud stained for the mRNA of tbxta (green in the posterior PSM and notochord), tbx16 (red in the PSM) and sox2 (blue in the spinal cord). **B.** Same image as in A. but only showing the sox2 (blue) channel. **C.** Same image as in A. but only showing the tbx16 (red) channel. **D.** Same image as in A. but only showing the tbxta (green) channel.

Once the tails had been staged and stained, they were imaged on a confocal microscope. The resulting images consist of four channels, one for each stained gene product plus DAPI, and we image over three spatial dimensions. Images were subsequently passed through a pre-processing pipeline in preparation to be used for network training and testing (see details in the Materials and Methods section). It is important to note that the tails in all images presented to the CNN were in the same orientation: anterior to the left, posterior to the right, dorsal up and ventral down as in Fig.1B and C. We obtained a total of 120 images stained with DAPI, 56 images stained for tbxta, 56 for sox2 and 56 for tbx16. Of the 120 DAPI stained images obtained, 96 were used for training (with 24 images per class) and the remaining 24 images were used for testing (with 6 images per class). Of the 56 images obtained for each gene of interest, 48 were used for training (with 12 images per class) and the remaining 8 were used for testing (with 2 images per class) (for more details please see the Final Data Sets section in the Materials and Methods and in particular Table 4). These are very small data set sizes compared to those usually used for deep learning classification problems, which typically range from the hundreds to the millions.

**Table 1.** Comparison matrix of classification outcomes of 2D and 3D morphological images. Training accuracy is derived from the highest average accuracy from all epochs and test accuracy, from testing on a subset of data that has not been used during training.

|  | DAPI |  |
| --- | --- | --- |
|  | Training accuracy (%) | Testing accuracy (%) |
| 2D | 100 | 100 |
| 3D | 100 | 79.16 |

**Table 2.** Classification outcomes of the CNNs trained with 3D gene expression image data. Training accuracy is derived from the highest average accuracy from all epochs and test accuracy, from testing on a subset of data that has not been seen during training.

|  | Ntl |  | Sox2 |  | Tbx16 |  |
| --- | --- | --- | --- | --- | --- | --- |
|  | Training accuracy (%) | Testing accuracy (%) | Training accuracy (%) | Testing accuracy (%) | Training accuracy (%) | Testing accuracy (%) |
| 3D | 95.55 | 100 | 100 | 100 | 100 | 87.5 |

**Table 3.** Classification outcomes of the CNNs trained with 3D gene expression image data. Training accuracy is derived from the highest average accuracy from all epochs and test accuracy, from testing on a subset of data that has not been seen during training.

|  | Ntl |  | Sox2 |  | Tbx16 |  |
| --- | --- | --- | --- | --- | --- | --- |
|  | Training accuracy (%) | Testing accuracy (%) | Training accuracy (%) | Testing accuracy (%) | Training accuracy (%) | Testing accuracy (%) |
| 2D | 72.77 | 75 | 80 | 62.5 | 79.44 | 87.5 |

**Table 4.** Data set summary. Our data were divided into two data sets, the 2D and the 3D datasets. Within those data sets there were image stained for DAPI, Tbxta, Tbx16 and Sox2. The table shows the total number of images of each kind (Total column), and how many images were used for training (Training column) and Testing (Testing column). In brackets, the number of images per class.

|  | 2D |  |  | 3D |  |  |
| --- | --- | --- | --- | --- | --- | --- |
|  | Total | Training | Testing | Total | Training | Testing |
| DAPI | 120 | 96 (24) | 24 (6) | 120 | 96 (24) | 24 (6) |
| Tbxta | 56 | 48 (12) | 8 (2) | 56 | 48 (12) | 8 (2) |
| Tbx16 | 56 | 48 (12) | 8 (2) | 56 | 48 (12) | 8 (2) |
| Sox2 | 56 | 48 (12) | 8 (2) | 56 | 48 (12) | 8 (2) |

All images are three dimensional, with *x, y* and *z* axes. By adding an an additional pre-processing step, we obtained a maximum intensity projection along the *z* axis for every image in order to obtain a two-dimensional representation of each image. In this way we generated an associated 2D version of the 3D data set. This dimensionality reduction was accompanied by a subsequent reduction in the size of the images compared with their 3D counterparts. As a result we were able to increase the dimensions of the *x* and *y* axes to 128 x 128 pixels. All data are available upon request.

### 0.2 Convolutional neural network architecture

We chose a simple architecture, composed of two convolutional layers, each using rectified linear unit (ReLU) activation functions followed by a max pooling operation (Fig 3). The first convolutional operation employs a kernel size of 5×5×5 and stride of 1×1×1 which produces 32 channels, and the second uses a kernel size of 4×4×4 and stride of 1×1×1, producing 64 channels. The number of channels produced by the convolution operation is arbitrary, and is often used as a hyperparameter to optimise a network. Both convolutions are followed by a max pooling layer with a kernel size of 3×3×3. Once passed through the convolutional layers, the data is flattened into a one dimensional array of length 262144 (64×64×64), and passed into the multi layered perceptron (MLP) part of the network. The MLP contains three fully-connected hidden layers, followed by a third output layer consisting of 4 units - one for each of our defined classes (16-18 somites, 20-22 somites, 24-26 somites and 28-30 somites). An activation function is implemented after each of the first two layers, and the dropout regularisation technique applied with a dropout rate of 0.2. The final layer of the network is a third fully connected layer containing four units which returns the log probabilities of each image belonging to each of the four classes. Finally, a softmax activation function is applied to these probabilities, which yields a percentage likelihood for each classification. During training, a cross entropy function is used as the loss function, which is optimised using the Adam optimisation function. The architecture described above corresponds to the 3D implementation of the classifier. The 2D implementation is generally identical, except for the necessary reduction in one dimension (Figs 3A and B, more details in the Materials and Methods section).

**Fig 3.**
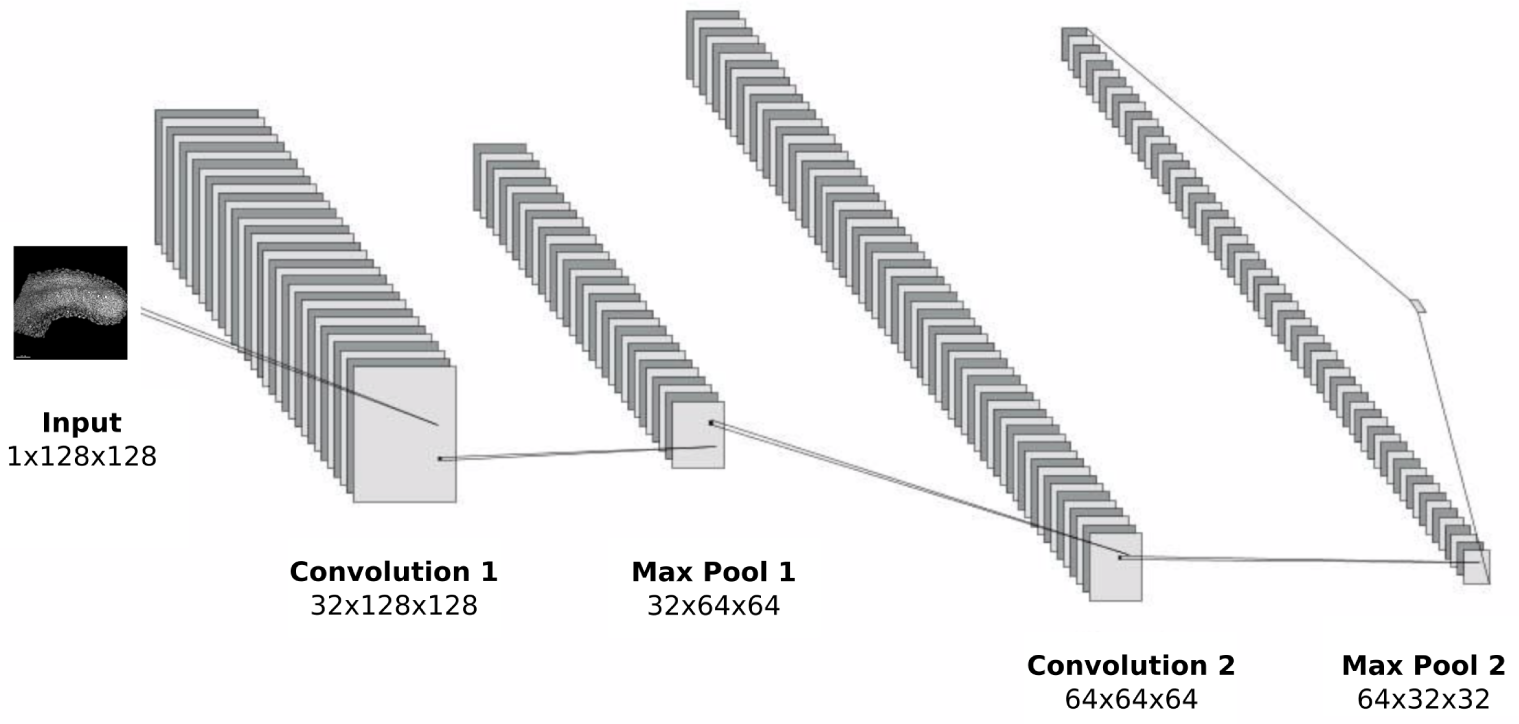
Convolutional neural network architecture. Convolution and pooling region of the 2D CNN architecture.

### 0.3 CNN-based classifiers trained on less than 100 morphological images of zebrafish tail buds reach up to 100% accuracy

Our first objective was to train a classifier that would be able to distinguish between morphological (DAPI stained) images of zebrafish tail buds at four discrete stages in development. The confocal images are three dimensional but can be reduced to two dimensions by performing a maximum intensity projection along one of the axes on the image analysis software Imaris, as detailed in the previous section. Given the small size of our data set, we wanted to see whether a 2D or a 3D data set would be most suitable to train a CNN to accurately classify zebrafish tail buds according to their developmental stage. To address this we trained both 2D and 3D versions of the network on 2D and 3D data sets respectively and compared them.

#### 0.3.1 CNN reaches close to 80% accuracy classifying 3D morphological images of zebrafish tail buds

The 3D CNN was initially parametrised using the optimisation algorithm SigOpt (see Materials and Methods section) to find the most appropriate activation functions and number of units in the first and second hidden layers. This procedure found that ReLU activation for both layers performed consistently better than Tanh activation for both layers, or a combination of the two. The algorithm settled on a first hidden layer composed of 501 units and a second composed of 397 units, which we rounded up to 500 and 400 units respectively (Materials and Methods, Table 5). All other network parameters remained as described in the previous section.

**Table 5.** Parameter optimisation results for the 3D CNN. The activation functions and hidden units refer to the parameters used in the first and second layers of the MLP part of the network, respectively. Test accuracy was obtained on a test data set of 24 images unseen by the network during training

| condition number | activation function 1 | activation function 2 | hidden units (layer 1, layer 2) | test accuracy (%) |
| --- | --- | --- | --- | --- |
| 1 | Tanh | Tanh | (664, 362) | 25 |
| 2 | Tanh | ReLU | (425, 391) | 29.16 |
| 3 | ReLU | Tanh | (694, 304) | 25 |
| 4 | ReLU | Tanh | (504, 497) | 29.16 |
| 5 | ReLU | ReLU | (473, 419) | 54.16 |
| 6 | ReLU | Tanh | (479, 347) | 20.83 |
| 7 | Tanh | ReLU | (564, 441) | 29.16 |
| 8 | Tanh | ReLU | (619, 336) | 33.33 |
| 9 | ReLU | ReLU | (501, 397) | 58.33 |
| 10 | Tanh | Tanh | (441, 421) | 25 |

This network was further trained on the 3D DAPI data set for 100 epochs to allow learning convergence and optimal classification results. The total training data set consisted of 96 3D DAPI stained images, with 24 per class (see Table 4). As expected, extending the training time results in a significant increase in accuracy and an associated decrease in loss. The training accuracy peaked at a perfect score of 100% after 100 epochs, compared to the 62% accuracy reached after 10 epochs during the initial parameter optimisation round. Similarly, test accuracy reached a maximum value of 79.16%, compared to the initial 58.33% test accuracy (Materials and Methods, Table 5). A test accuracy of almost 80% is already a very good score, and a surprising one too, considering the small size of the training set used.

#### 0.3.2 CNN reaches 100% accuracy classifying 2D morphological images of zebrafish tail buds

Next, we proceeded to train the 2D version of the classifier on the associated 2D data set. As with the 3D classifier, we use an initial course grained parameter optimisation strategy to find the best performing parameter combination, before fine training the selected network for longer times (more epochs, see Materials and Methods). The resulting network again uses ReLu activation functions, this time with 429 and 330 units in the first and second hidden layers respectively (see Table 6, cyan). Using these parameters, the network converged after only 25 epochs. This is a small number of epochs compared to the 100 that were required for convergence in the 3D classifier. The resulting accuracy on the test data set reached a perfect accuracy of 100%. This was expected since since in the course-grained training this network had already scored 95.83% (see Table 6, cyan row represents the parameters that result in the best performance).

**Table 6.** Parameter optimisation results for the 2D CNN. Hidden units refer to the parameters used in the first and second layers of the MLP part of the network respectively. Test accuracy was obtained on a test data set of 24 images unseen by the network during training

| condition number | hidden units layer 1 | hidden units layer 2 | test accuracy (%) |
| --- | --- | --- | --- |
| 1 | 452 | 431 | 83.33 |
| 2 | 691 | 480 | 91.66 |
| 3 | 428 | 323 | 83.33 |
| 4 | 458 | 398 | 91.66 |
| 5 | 504 | 463 | 87.50 |
| 6 | 587 | 438 | 87.50 |
| 7 | 418 | 327 | 95.83 |
| 8 | 609 | 381 | 87.50 |
| 9 | 654 | 320 | 91.66 |
| 10 | 428 | 429 | 91.66 |

#### 0.3.3 Performance comparison of 2D vs 3D CNN-based classifiers of morphological images of zebrafish tail buds

Both classifiers perform exceptionally well, especially considering that less than 100 images were used to train four different classes. The 3D classifier reached an accuracy of almost 80% while the 2D classifier achieved an accuracy of 100% (as shown in Table 1). We had initially expected the increased information contained in the 3D images when compared to their 2D counterparts, to result in a better performance of the 3D classifier.

Instead we find the opposite. Furthermore, the 2D network obtained a higher test accuracy at a quarter of the computational time, converging after only 25 epochs as opposed to the 100 required to converge the 3D network. One possible explanation for this result might have to do with our methodology: to train the 2D network we initialise using the some parameter values obtained from training the 3D network in a somewhat unusual application of transfer learning. Transfer learning refers to using the weights obtained by training a network with a given data set, as the starting point for training the network on a different data set and for another classification task [58]. This method has already been shown to be extremely effective at reducing the amount of labelled image data required for training CNNs on biological microscopy data [25, 41, 59–61]. Usually this approach is used to train networks on new data sets and classification challenges. In our case, transfer learning was applied to the same classification task and data, albeit in a different format, which yielded an improved performance and faster convergence time.

In practical terms, this result dispels the myth that training a CNN to a high test accuracy always requires prohibitive amounts of data, and shows how transfer learning can help make the most out of the available data, in this case to further improve the levels of accuracy to obtain a perfect score on the 2D version of the same set. This result suggests that CNNs can be successfully applied to myriads of classification tasks in the life sciences, where data are scarce.

### 0.4 CNN-based classifiers trained on less than 50 3D images of gene expression domains in zebrafish tail buds reach up to 100% test accuracy

Next, we applied a traditional transfer learning approach to ask whether 2D and 3D classifiers could be trained to stage gene expression domains on zebrafish tail buds at the same four developmental stages considered before (Fig.2B, C and D). To do this, we used the network architecture and weights obtained in the previous section from training using 2D and 3D morphological (DAPI stained) images of zebrafish tail buds, and re-trained these networks using the gene expression data. Gene expression data is fundamentally different from the previously used DAPI stains (compare Fig.1A to Fig.1B and Fig.2B, C and D). While DAPI stains the nucleus of every cell, hence building an image of the tail bud’s morphology, HCR V.3. stains the mRNA of the gene of interest anywhere in the cell and surroundings, resulting in hazy coloured clouds. To add to the challenge of staging hazy gene expression domains, the data sets used were half the size with respect to those used to train for morphological classification in the previous section (Table 4), with a training set size of 48 images for each gene.

#### 0.4.1 CNNs stage 3D gene expression domains in zebrafish tail buds with a minimum of 87.5% accuracy

We used the 3D network obtained by training using the 3D morphological image data set (previous section) as the starting point from which to train three networks, each of which will learn to classify 3D gene expression image data for one gene (Sox2, Tbx16 or Ntl respectively. See Fig 2). The gene expression image data set for each gene is smaller than the morphological image data sets used in the previous section, with each containing less than half the number of samples (56 as opposed to 120, see Materials and Methods, Table 4). To reduce the risk of over-fitting that is commonplace when training networks with small data sets, we reduced the number of epochs to 25. Preliminary experiments showed that the models tended to converge quicker with the smaller data sets, corroborating our choice of a reduced training time.

After 25 epochs, the networks trained to stage 3D Ntl and Sox2 expression domains in the tail bud both achieved a test accuracy of 100% while the network trained to stage 3D Tbx16 expression patterns reached a lower, but also impressive, maximum test accuracy of 87.5% (Table 2). The observed differences in accuracy could be due to a number of reasons including differences in the level of background staining between each of the three genes or they could be rooted in the existing differences between the shapes of the expression domains themselves. The rates of convergence during training did not differ significantly between the three networks (S1 Fig).

#### 0.4.2 CNN-based classifiers trained on less than 50 2D images of gene expression domains in zebrafish tail buds score between 60 and 90% test accuracy

Again, we use a transfer learning approach where the 2D network pre-trained on morphological image data (section 0.3) was taken as the starting point from which to train three new networks, each of which will learn to stage 2D gene expression domains. As in the 3D case discussed in the previous section, we train for 25 epochs to reduce potential over-fitting.

For each gene, the training accuracy of the 2D models plateaus soon after 10 epochs achieving between 70% and 80% in each case (Table 3). The range of testing accuracy that we obtain for the 2D expression data is larger than that obtained from training: between 60% and 90% (Table 3) and performing overall much worse than when the networks were trained on 3D expression data.

#### 0.4.3 Performance comparison of 2D vs 3D CNN-based classifiers of gene expression images

Contrary to what we found with the networks trained on morphological (DAPI-stained) tail bud images, networks trained on gene expression data sets tended to perform consistently better when trained on the 3D image data as opposed to those trained on 2D image data (except for Tbx16 where the test accuracy of the 3D and 2D networks are the same). It is difficult to pinpoint exactly the reasons underlying the improved performance of the 3D network, however it is possible that for these very small data set sizes, the training favours from the extra information contained in the 3D images, and is able to take full advantage of it. The 2D images are rendered by projecting down all of the information in the *Z* axis, hence averaging it out. While it seems that this information is expendable when training on DAPI-stained images, it is definitely not when training on gene expression data. This suggests, at least for such small training set sizes, that the changes undergone by gene expression domains in all dimensions are learnt and used by the network.

## Conclusion

Our results have shown that two and three dimensional convolutional neural networks can be trained to stage developing zebrafish tail buds based on both morphological and gene expression images. Importantly, we show that high accuracies can be achieved with data set sizes of under 100 images, much smaller than the typical training set size for a convolutional neural net, which tend to be at least in the tens of thousands range, if not larger.

By showing that we can we build a CNN-based classifier to stage isolated structures such as tail buds, our work also highlights that it is not necessary to always rely on established developmental landmarks such as somite formation for staging. This ability will make it possible to directly stage developmental processes that are difficult to otherwise stage accurately based solely on such landmarks. Furthermore, image series can be generated directly from tissue explants to train a CNN-based classifier, leaving out entirely the additional step of having to stage whole embryos. This approach should prove particularly useful considering the recent rise in the use of 3D culture methods to derive specific tissue derivatives, or organoids. Research on these systems requires the development of accurate staging systems since a tissue or organ of interest developed in isolation on a dish can no longer be staged relative to the development of the whole embryo [62]. A recent example highighting the importance of being able to stage such structures comes from work were the ability to order intestinal organoids along a common morphogenetic trajectory was key for determining the mechanisms of symmetry breaking at the early stages of their development [63]. We propose that the development of CNN-staging methods will offer a broad range of advantages, allowing us to follow developmental events in both embryonic explants and organoids.

Convolutional neural nets are becoming increasingly wide-spread and thanks to the various user-friendly tools that are now available to implement them, we expect that their use to soon extend even further. Although our results suggest that the data requirements for training a CNN-based classifier are highly dependent on the nature of the data themselves, this work constitutes a proof of principle that we hope will contribute to dispel the myth that large data set sizes are always required to train CNNs, and encourage researchers in fields where data are scarce to apply ML approaches.

## Materials and methods

### Ethics statement

This research was regulated under the Animals (Scientific Procedures) Act 1986 Amendment Regulations 2012 following ethical review by the University of Cambridge Animal Welfare and Ethical Review Body (AWERB).

### Zebrafish husbandry, manipulation and embryo collection

Zebrafish from the wild-type Tupfel Long-fin line zebrafish (Danio rerio) were kept at 28.5°C, as recommended by standard protocols. Crosses with one male and one female were set up and left overnight. The morning after, dividers were lifted and the fish allowed to mate for 15 minutes, after which the embryos were collected. This was done to favour the synchronous developmental of embryos in the same batch. Embryos are kept in E3 embryo medium (Westerfield, 2000) and incubated at between 26°C and 32°C until they reach the correct somite stage range of one of the four classes (16-18 somites, 20-22 somites, 24-26 somites and 28-30 somites) according to the Kimmel *et al* (1995) staging table. Embryos are de-chorionated by hand, fixed in 4% paraformaldehyde (PFA) and stored overnight at 4°C.

### *In situ* hybridization

In situ hybridization was performed using third-generation DNA hybridization chain reaction (HCR V3.0), and carried out as described by Choi *et al.* (2018). In brief, embryos are incubated overnight at 37°C in 30% probe hybridisation buffer containing 2pmol of each probe mixture. Excess probes were washed off with 30% probe wash buffer at 37°C and 5xSSCT at room temperature. Embryos are then incubated overnight in the dark, at room temperature in amplification buffer containing 15pmol of each fluorescently labelled hairpin. A shaker was used to ensure a thorough mixing of the hairpin solution, which allowed us to increase the number of embryos per eppendorf approximately 10-fold. Following HCR, embryos were bathed in DAPI overnight at 4°C. The protocol takes a total of three days. Probe sequences were designed and manufactured by Molecular Instruments, Inc.

### Sample preparation and imaging

A precision scalpel was used to dissect the posterior unsegmented and tail bud regions from the embryo bodies while viewing through a Nikon Eclipse E200 in bright field at 10x objective. Tail buds were then collected using a glass pippette, placed in the center of a glass-bottom dish and mounted using methyl-cellulose. Particular emphasis was put on trying to not make the cuts stereotypical, in order to reduce the likelihood of them being learnt by the neural net. Images were acquired using a Zeiss LSM 700 laser-scanning confocal microscope, and the accompanying Zeiss Zen 6.0 software. The z-dimension and laser intensity settings were modified on a batch-by-batch basis to account for variance in the depth range of the samples between experiments, while the bit depth and pixel dimensions for the image capture were kept constant at 12-bit and 256 x 256px, respectively.

### Image processing

Image samples were processed using Imaris 9.2.2 (Bitplane) and FIJI (Schindelin *et al.*, 2012). Each image was processed manually according to the following workflow:

1. Images are first saved in Zeiss’ proprietary LSM (.lsm) format and are then imported into Imaris. Each image is repositioned using the ‘Image Processing -¿ Free Rotate’ function. The tail bud is moved in three dimensions such that the antero-posterior (AP) and dorso-ventral (DV) axes are approximately positioned left-to-right and top-to-bottom, respectively.
2. A surface is created using the DAPI channel. This surface is used to segment the whole region of the image taken up by the unsegmented posterior embryonic axis and tail bud. Masks are then used to set all voxels outside of this generated segment to zero on all four channels, leaving the target region intact while removing all imaging noise/artifacts from the sample. 3D images of each of the four channels (DAPI, Tbxta, Sox 2 and Tbx16) were exported individually in .TIF format. In addition, maximum intensity projection images (projections onto the z axis) were also obtained and exported for each channels and used to assemble the 2D datasets (details below).
3. Once in .TIF format and before presenting the images to the model for training, they are subject to a simple programmatic pre-processing pipeline where, first, the *z* axis is normalised for all training samples. As a result, images which initially had varying counts of z-slices due to the batch-by-batch nature of the image acquisition procedure, will now all have been normalised to the same size, as required by the neural net. Next, it is necessary to crop the *x* and *y* dimensions of each slice to accommodate the computational bottleneck of fitting heavy 3D images into memory. The last pre-processing step before inputting the data into the network requires us to convert the images into Numpy array format (Walt *et al*, 2011). Numpy arrays are an efficient and robust way of representing image data in the form of a multi-dimensional matrix, and is how the downstream model will process and understand the contents of the image.
4. Finally, images were converted into grayscale and reshaped to 1 x 64 x 64 x 64 pixels, where the first dimension represents the number of channels (1 for a grayscale image) before being presented to the network.

### Final Datasets

The final data sets consists of 2D and 3D collections of images for DAPI stained embryos, as well as embryos for stained using HCR V3.0 for the gene products of three genes: Tbxta, Sox2 and Tbx16. The 2D and 3D DAPI image datasets contain each a total of 120 images: 96 of which are used for training - 24 images for each class - and 24 images for testing - 6 per class. The 2D and 3D HCR image datasets contain a total of 56 images for each gene: in each case, 48 are used for training (12 per class), and 8 were used for testing (2 for each class). An equal split of samples per class maintained in all cases. No data augmentation was used.

### 2D CNN formulation

The CNN presented in the Results section corresponds to the 3D implementation of the network. The 2D implementation is generally identical, except for the necessary reduction of one dimension. To achieve this, the kernel size for the first convolution layer is reduced from 5×5×5 (in the 3D model), to 5×5. The flattened layer must also be adapted to the reduction in dimensions, and in the 2D implementation takes the form of a vector of length 65536. Figs 3A and B show a visual representation of the network architecture for the 2D network. All other parameters were kept the same as in the 3D network.

### Network training and initial parameter optimisation

The parameter optimisation algorithm SigOpt was used to parametrise the 3D classifier. Given a series of parameter options, this algorithm will run the network with different parameter combinations as it optimises the performance of a previously defined target metric. The metric used in this case was the test accuracy, that is the accuracy of the classifier when applied to a subset of images that have not been seen during training. We allowed each parameter set to be trained during only 10 epochs. This suffices to highlight the best performing networks while economising the time spent training the network.

The parameters to be optimised corresponded to the type of activation functions to be used within each of the first two layers of the MLP region of the network, and the number of hidden units in those same layers. Activation functions being considered were ReLU and Tanh, and the search range for optimal number of units was 400-700 units for the first hidden layer and 300-500 for the second.

The outcome of the optimisation revealed that for this task the ReLU activation function was consistently superior to Tanh, with the highest test accuracy scores coming from the networks which had a ReLU activation function at both layers (Table 5, conditions 5 and 9, gray and cyan rows). When comparing these conditions we realise that the precise number of hidden units in each layer seems to have less influence on the resulting test accuracy. We chose parameters in condition 9, rounded to nearest 10 hidden units, as these gave the highest accuracy: 58.33% and re-trained this network with a significantly increased training time (100 or 20 epochs depending on the data, see Results section) to evaluate its maximum performance.

As with the 3D network, we use an initial course grained parameter optimisation strategy using the SigOpt algorithm to find the best performing parameter combination of the 2D network before fine training the selected network for longer times. We decided to use ReLu activation functions in order to keep the 3D and 2D networks as similar and therefore comparable to each other as possible. Instead, only the number of units in the hidden layers were optimised.

The optimisation process yielded a best configuration with a hidden unit count of 420 and 330 for the first and second layers, respectively (Table 6, condition 7, cyan row).

### Code and computer specifications

The code for this project was written in Python (Python Software Foundation), making use of the large ecosystem of data manipulation tools and libraries available therein.

The machine learning model development was set up using Pytorch, a Python API of the Torch ML framework. Pytorch allows for performing operations on a graphics processing unit (GPU), which parallelizes the large mathematical operations, and increases performance time significantly. The GPU used in this project was an Nvidia 2080Ti RTX. All code used in this project is available at: https://github.com/ajrp2/ssclassifier

## Supporting information

**S1 Fig.**
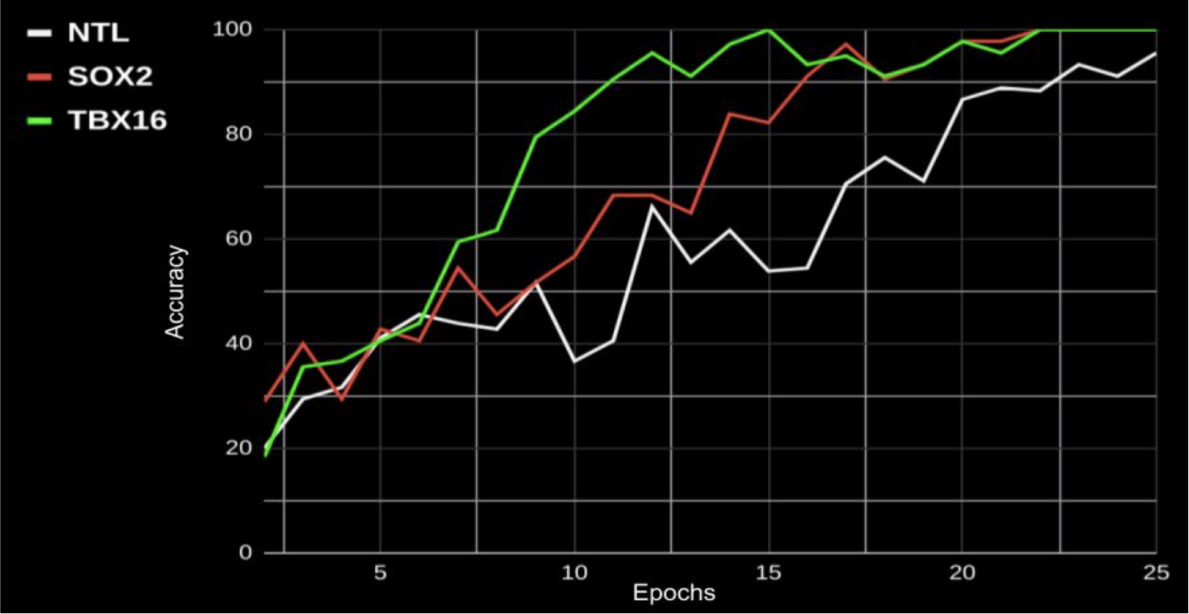
Training accuracy of the 3D CNNs when trained on 3D gene expression data. Accuracy is obtained at every epoch by averaging the accuracy scores for that particular epoch.

**S2 Fig.**
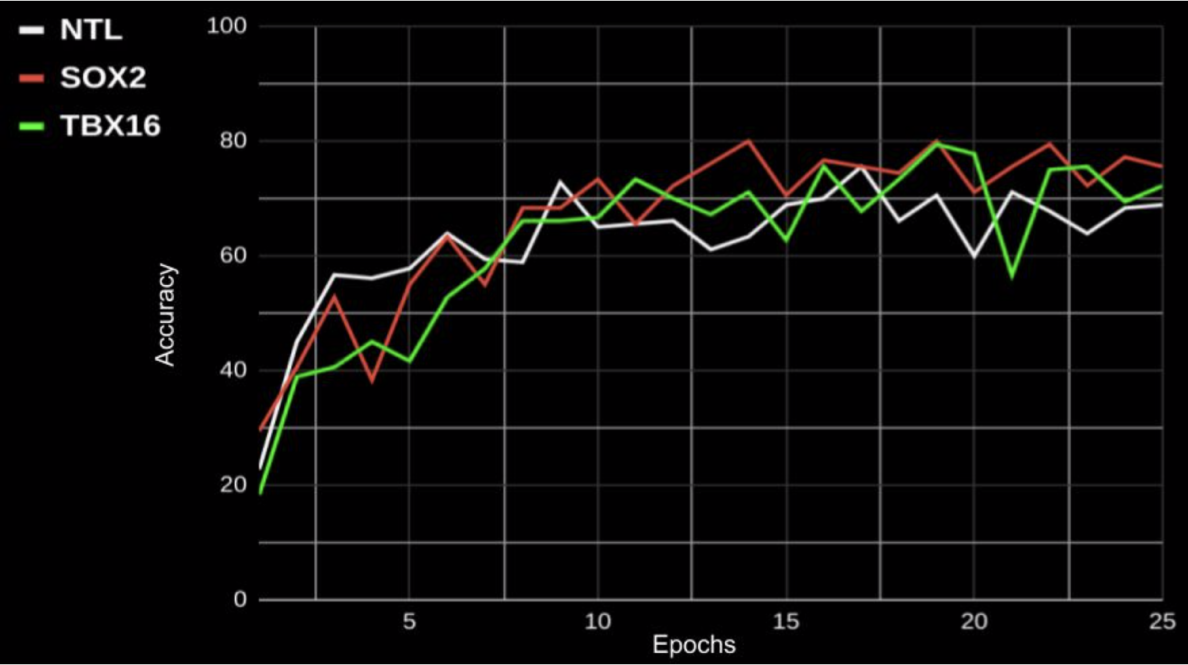
Training accuracy of the 2D CNNs when trained on 2D gene expression data. Accuracy is obtained at every epoch by averaging the accuracy scores for that particular epoch.

## Acknowledgments

The authors would like to thank Rubén Pérez-Carrasco and Elia Benito-Gutiérrez for their helpful comments on and extended version of this manuscript B.S. and S.H were supported by a Henry Dale Fellowship jointly funded by the Wellcome Trust and the Royal Society (109408/Z/15/Z). B.V. was funded by a Herchel Smith Fund Postdoctoral Fellowship.

